# Sex-specific DNA methylation differences in Amyotrophic lateral sclerosis

**DOI:** 10.1101/2024.11.22.624866

**Authors:** Olivia A. Grant, Alfredo Iacoangeli, Ramona A. J. Zwamborn, Wouter van Rheenen, Ross Byrne, Kristel R. Van Eijk, Kevin Kenna, Joke J.F.A. van Vugt, Johnathan Cooper-Knock, Brendan Kenna, Atay Vural, Simon Topp, Yolanda Campos, Markus Weber, Bradley Smith, Richard Dobson, Michael A. van Es, Patrick Vourc’h, Philippe Corcia, Mamede de Carvalho, Marc Gotkine, Monica P. Panades, Jesus S. Mora, Jonathan Mill, Fleur Garton, Allan McRae, Naomi R. Wray, Pamela J. Shaw, John E. Landers, Jonathan D. Glass, Christopher E. Shaw, Nazli Basak, Orla Hardiman, Philip Van Damme, Russell L. McLaughlin, Leonard H. van den Berg, Jan H. Veldink, Ammar Al-Chalabi, Ahmad Al Khleifat

## Abstract

Sex is an important covariate in all genetic and epigenetic research due to its role in the incidence, progression and outcome of many phenotypic characteristics and human diseases. Amyotrophic lateral sclerosis (ALS) is a motor neuron disease with a sex bias towards higher incidence in males. Here, we report for the first time a blood-based epigenome-wide association study meta-analysis in 9274 individuals after stringent quality control (5529 males and 3975 females). We identified a total of 226 ALS saDMPs (sex-associated DMPs) annotated to a total of 159 unique genes. These ALS saDMPs were depleted at transposable elements yet significantly enriched at enhancers and slightly enriched at 3’UTRs. These ALS saDMPs were enriched for transcription factor motifs such as ESR1 and REST. Moreover, we identified an additional 10 genes associated with ALS saDMPs through chromatin loop interactions, suggesting a potential regulatory role for these saDMPs on distant genes. Furthermore, we investigated the relationship between DNA methylation at specific CpG sites and overall survival in ALS using Cox proportional hazards models. We identified two ALS saDMPs, cg14380013 and cg06729676, that showed significant associations with survival. Overall, our study reports a reliable catalogue of sex-associated ALS saDMPs in ALS and elucidates several characteristics of these sites using a large-scale dataset. This resource will benefit future studies aiming to investigate the role of sex in the incidence, progression and risk for ALS.

## Introduction

Amyotrophic lateral sclerosis (ALS) is a debilitating neurodegenerative disease characterized by the progressive loss of motor neurons in the brain and spinal cord^1^, leading to muscle weakness, paralysis, and ultimately respiratory failure. Its impact extends beyond the individual, affecting their families and straining healthcare systems worldwide. Despite its prevalence, the exact causes of ALS remain elusive. Moreover, the cost of this disease to healthcare systems also remains a growing issue for several economies across the world, adding to the increasing pressure to pursue further research in this area of science.

Genetic factors play a significant role in ALS with approximately 10–15% of ALS cases being considered familial cases^2^. Familial ALS is inherited in either an autosomal dominant, autosomal recessive, or X-linked mode, while the remaining are sporadic (SALS) meaning the exact cause is not linked to inherited genetic mutations^3^. About 70% of FALS and 15% of SALS have mutations in known ALS genes, including *SOD1, FUS, TARDBP, C9ORF72, ATXN2*, and more^4^. These genes are mainly implicated in protein homeostasis, RNA metabolism, vesicle transport, mitochondrial function, and more.

However, a substantial proportion of ALS cases lack identifiable genetic mutations, suggesting that environmental and lifestyle factors may also play a crucial role. Numerous studies have associated smoking and exposure to certain toxins with an increased risk of developing ALS^5–9^. These findings underscore the complex interplay between genetic susceptibility and environmental influences in ALS pathogenesis.

In recent years, there is an increasing interest into which role epigenetic mechanisms may play in both gene-environment interactions and the broader impact of the environment on gene expression. Epigenetics refers to changes in gene expression that do not involve alterations in the DNA sequence itself^10^. Three main epigenetic marks may play pivotal roles: histone modifications, RNAs, and DNA methylation^10^. DNA methylation, one of the most studied epigenetic modifications and the mark we will focus on in this research, involves the addition of a methyl group to a cytosine base and can regulate gene activity and reflect past environmental exposures. Therefore, understanding epigenetic changes in ALS can provide insights into the mechanisms underlying disease development and progression^11^.

To this effect, past research studies have focused on deciphering the DNA methylation signatures related to ALS^12–18^. The most sizeable study in 2022 performed a blood-based epigenome-wide association study meta-analysis in 9706 individuals and identified a total of 45 differentially methylated positions (DMPs) which were annotated to genes enriched for pathways and traits related to metabolism, cholesterol biosynthesis, and immunity^12^. Further, they demonstrated that DNA methylation at numerous DMPs was associated with survival rate in individuals with ALS, suggesting they may have potential as biomarkers for predicting patient outcomes and guiding personalized treatment strategies. Another study noted increased global DNA methylation levels in the blood of ALS cases when compared to controls which was further correlated with disease duration too^17^. Notably, this finding is contradicted by research conducted using twin pairs that suggests that there is no trend associating genomic methylation with disease status^18^. However, they and other research groups have identified several DMPs showing differential methylation related to disease status^18, 16^, indicating that individual loci may be more informative with regards to investigating the interplay between epigenetics and ALS.

Despite the substantial progress in identifying DNA methylation signatures related to ALS, significant gaps remain in our understanding of how these changes may differ between sexes. It is well-documented that ALS exhibits sex-specific characteristics, for example, men are more commonly affected by ALS, with a ratio typically exceeding 1.5:1, suggesting a higher susceptibility among males^19^. Moreover, men tend to develop ALS at a younger age, typically in their late 50s to early 60s, compared to women who are often diagnosed older^20^. Moreover, studies hint at a potential difference in disease progression rates between the sexes due to exposure to different levels of sex hormones^21^. These variations underscore the influence of sex-specific genetic and environmental factors on ALS manifestation and progression.

Understanding whether DNA methylation changes in ALS are sex-specific is crucial for several reasons. First, it could elucidate the underlying mechanisms that contribute to the observed differences in disease manifestation and progression between men and women. Second, it may reveal novel biomarkers for ALS that are specific to each sex, thereby improving diagnostic accuracy and prognostic predictions. Lastly, this knowledge could inform the development of sex-specific therapeutic strategies, potentially enhancing the effectiveness of treatments.

Our study aims to address this gap by conducting a meta-analysis of sex differences in ALS using DNA methylation data from 9274 individuals provided by Project MinE. By examining epigenetic patterns across sexes, we seek to uncover potential biomarkers and pathways implicated in ALS pathogenesis. This approach will not only advance our understanding of the sex differences present in this complex disease but also pave the way for more personalized and effective interventions.

## Results

### Several known ALS loci show sex differences in DNA methylation

Our sex-specific meta-analysis was performed using linear regression on DNA methylation (DNAm) data obtained from the Illumina 450k BeadChip and EPIC BeadChip platforms, encompassing 9,274 individuals. This included 3,975 females and 5,299 males. We excluded known SNP probes, cross-hybridizing probes, and X/Y-linked probes from the analysis. Further, given that whole blood is a bulk tissue, we estimated the cell type proportions between male and female individuals to determine whether any differences in these proportions could introduce false positives in our results.

To ensure that differences in whole blood cell composition by sex would not bias our findings, we conducted a comparative analysis of the estimated cell type proportions between male and female individuals. Our analysis showed no significant differences in the relative proportions of the assessed cell types between sexes, suggesting that variations in cell composition were unlikely to confound our results.

However, when investigating sex-specific differences in cell type proportions in the context of ALS, we found distinct patterns of immune cell composition between ALS cases and controls, with notable differences between males and females. Specifically, CD4+ T cells exhibited a statistically significant sex difference unique to ALS cases, as determined by the Wilcoxon rank-sum test. Female ALS cases showed higher proportions of CD4+ T cells compared to their male counterparts, a difference not observed in the control group (Supplementary Figure 1). These findings suggest a potential sex-specific immune response involvement in ALS, possibly influencing disease progression and outcomes. While no significant sex-based differences in cell type proportions were found in the general population, the observed variation in CD4+ T cells within the ALS cohort highlights the importance of considering sex as a biological variable in ALS research.

As a result, we included cell type proportions in our models for identifying sex-associated differentially methylated probes in our epigenome-wide association studies. Additionally, the mean age at blood collection was 62.670 years for females and 62.37 years for males. Finally, we adjusted for test inflation using bacon since the lambda values were slightly high (Supplementary Figure 2).

After adjusting for multiple testing using the Benjamini–Hochberg FDR method (adj p value <0.05), we identified 226 autosomal CpGs that are associated with ALS and also display a significant sex difference in DNA methylation (Figure 1A).

**Figure 1.**
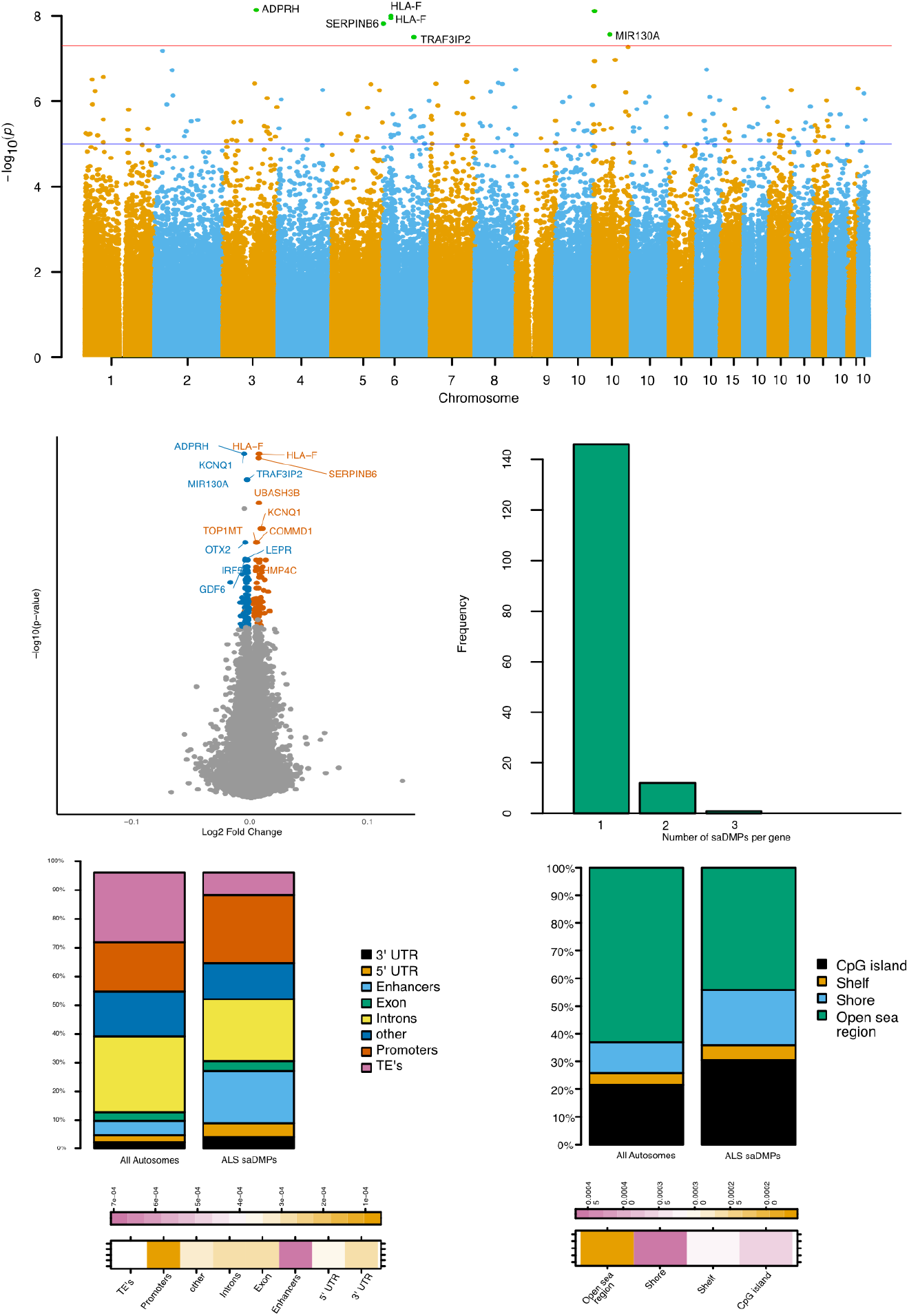
Location and characterisation of ALS saDMPs. A) Manhattan plot for meta analysis of sex associated ALS DNA methylation signatures. CpG sites which met a threshold of FDR<0.05 in meta analysis are represented above the blue line. CpG sites which met a threshold of Bonferonni < 0.05. B) Volcano plot for saDMPs. CpGs which are not significant in meta analyses are represented in grey, male-biased ALS saDMPs are in orange and female-biased ALS saDMPs in blue. C) Stacked barplots representing the overlap of ALS saDMPs with genomic features compared to the autosomal background. D) Heatmap showing the log2 (obs/exp) annotations based on the autosomal background of the different annotations.

**Figure 2.**
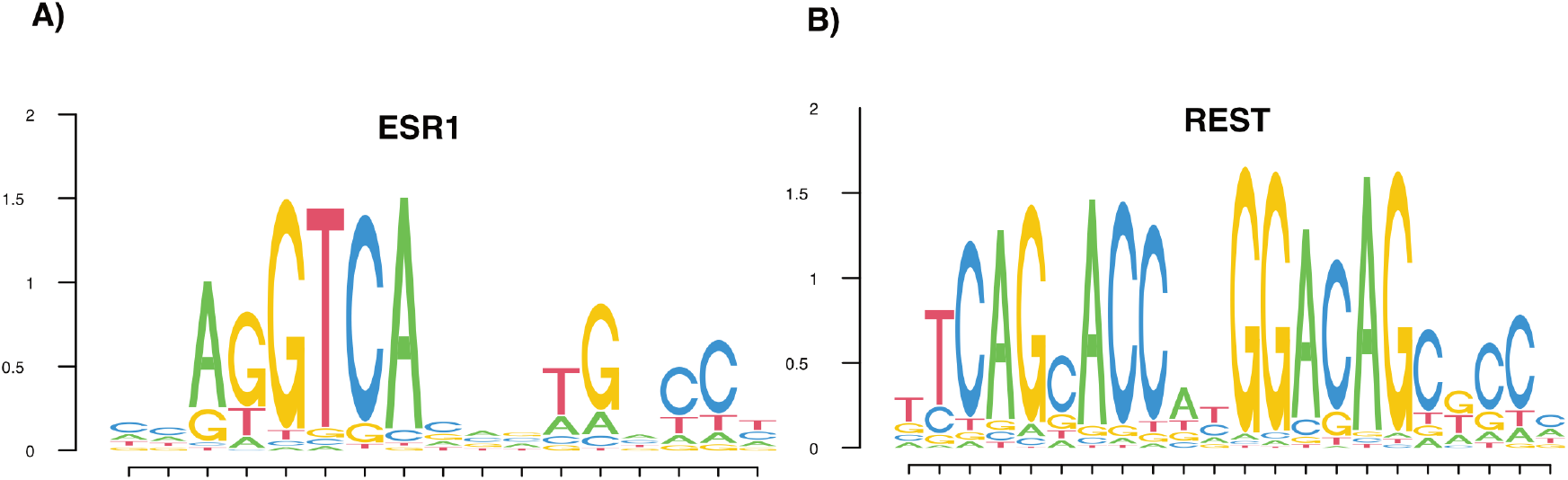
Enriched DNA binding motifs for ESR1 and REST, which were identified as enriched transcription factor motifs among the ALS-associated saDMPs.(A) The DNA binding motif for ESR1. (B) The DNA binding motif for REST. The height of each letter in the sequence logo represents the relative frequency of each nucleotide at that position, indicating the specificity and preference of the transcription factor for each base in the motif.

Despite identifying a significant number of CpGs associated with ALS that also display a significant sex difference in DNA methylation, the actual differences in methylation levels between sexes are small. Using an FDR threshold allowed us to capture these subtle changes while maintaining a rigorous approach to significance, although alternative methods like permutation thresholds could potentially reduce the number of significant findings as highlighted by the bonferroni threshold identifying a conservative 7 ALS saDMPs (Figure 1A). Of the 226 autosomal CpGs identified, 54% were more highly methylated in females (female biased), and the remaining 46% were more methylated in males (male biased). While these findings highlight a statistically significant sex difference, the biological magnitude of these differences is subtle.

This subtle yet statistically significant difference suggests that while sex-specific methylation patterns exist, the overall impact of these differences on gene expression and disease mechanisms might be limited. However, although the observed methylation differences between sexes were relatively small, these subtle changes can be biologically significant, particularly if they occur in regulatory regions that control the expression of genes crucial for neuronal health. Accumulation of these minor changes over time could potentially contribute to the onset and progression of ALS.

We performed pathway analysis using both Gene Ontology (GO) and KEGG databases, but no significant pathways were identified. The lack of significant enriched terms in GO and KEGG pathway analyses further supports the notion that these sex-specific methylation differences do not prominently cluster within particular biological pathways or processes. This diffuse pattern of methylation changes may imply a more complex interplay of genetic and epigenetic factors in ALS, where the sex differences in methylation are part of a larger, multifactorial context influencing disease development and progression.

### Characterisation of ALS saDMPs

The saDMPs were found in 159 unique genes with 13 of these genes harbouring several saDMPs including *DNAJC6, FLRT2, HLA-F* and more (Figure 1C). The number of saDMPs harboured by individual genes ranged from 1 to 3. HLA-F, a gene known to be display genetic sex differences in ALS cases also harboured the largest number of ALS saDMPs, 3, which interestingly were all found to be male-biased CpGs. Further, these CpGs were all located in the CpG shore region of the gene, an area known to be typically unmethylated and influence gene expression^22^. The top 10 significant CpGs include CpGs located several known ALS genes such as *KCNQ1, HLA-F, MIR130A*^23–25^ and also some novel ALS loci such as *ADPHR* (Table 2). Further, we highlight that the top 7 CpGs were not only FDR significant but also Bonferroni significant, underscoring these as strong associations. To gain a better understanding into a potential functional role of these ALS SADMPs, we investigated the genomic location of these ALS saDMPs in two different ways. Firstly, we looked at their annotations relative to CpG islands, compared to the EPIC background. Through this we were able to identify that the ALS saDMPs were in fact enriched in CpG shores, slightly enriched at CpG islands and interestingly, depleted at open sea regions of the genome. Due to the relationship between CpG island and CpG shore methylation with gene regulation, this suggested these ALS saDMPs could potentially have a functional role. To explore this idea further, we additionally annotated these ALS saDMPs to additional regions such as 3’UTRs, 5’UTRs, enhancers, exons, introns, promoters and transposable elements. Here, we identified that these ALS saDMPs were significantly depleted in transposable elements compared to the EPIC background, however interestingly these ALS saDMPs are slightly enriched at 5’UTRs and significantly enriched at enhancers suggesting that these sites may play a role in gene regulation.

**Table 1.** Characteristics of the study population across different datasets, including disease status, gender distribution, average age (in years), and survival status.

|  | 450k | EPIC | AUS 1 | AUS 2 |
| --- | --- | --- | --- | --- |
| **Disease status** |  |  |  |  |
| Control | 1428 | 756 | 595 | 148 |
| Case | 3006 | 2749 | 493 | 99 |
| **Sex** |  |  |  |  |
| Females | 1846 | 1518 | 487 | 124 |
| Males | 2588 | 1987 | 601 | 123 |
| **Average age (years)** |  |  |  |  |
| Females | 62.70 | 61.29 | 70.21 | 60.89 |
| Males | 62.42 | 60.97 | 70.42 | 59.49 |
| **Survival status** |  |  |  |  |
| Alive | 1748 | 1645 | 0 | 0 |
| Dead | 2527 | 1679 | 0 | 0 |
| Missing | 0 | 0 | 1088 | 247 |

**Table 2.**
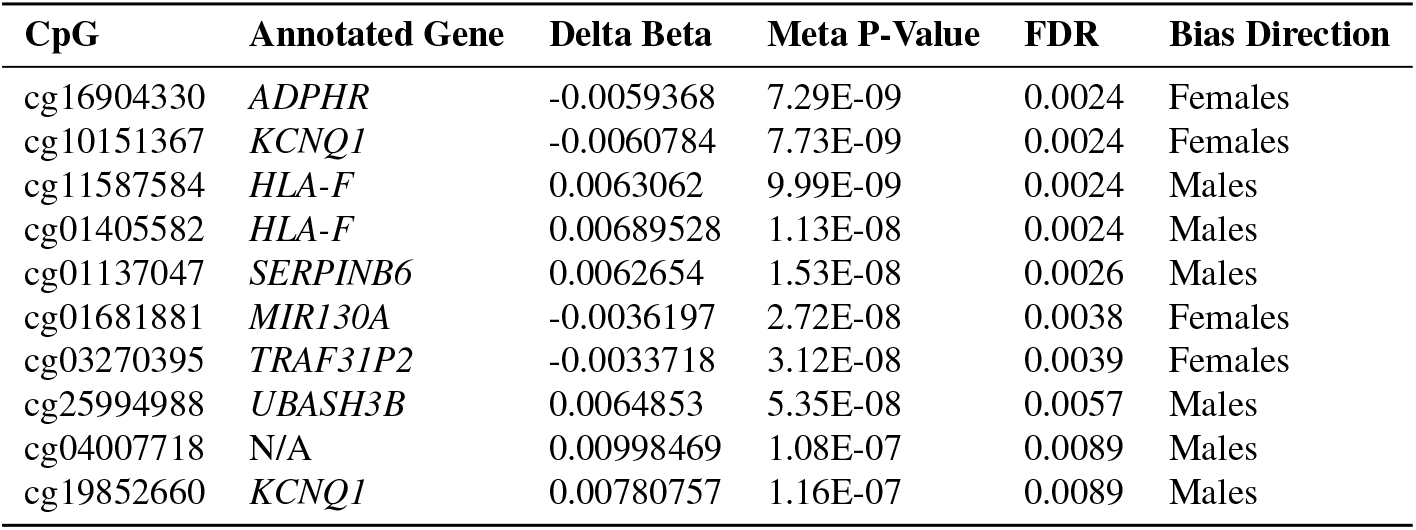
Top 10 significant ALS sex-associated differentially methylated positions identified by meta-analysis of 4 cohorts.

The presence of saDMPs at enhancers suggests that these saDMPs may potentially be involved in regulating distant genes. We therefore investigated the possibility that these ALS saDMPs may contact the promoter regions of known ALS genes through chromatin loops. Chromatin loops are 3D structures formed when distant regions of the genome are brought into close spatial proximity through physical interactions. This looping mechanism helps organize the genome within the nucleus and is thought to play a role in regulating gene expression. Through this investigation, we were able to annotate the ALS saDMPs to a further 10 genes (Table 3). For example, a female biased ALS saDMP located on chromosome 5 annotated to gene (LINC00461) with no previously reported direct link to ALS itself was found to potentially contact the promoter region of the *SLC36A1*gene, which has previously been reported to carry a genetic risk for ALS and Alzheimers. Further, another female biased ALS saDMP on chromosome 17 annotated to gene *CAVIN1* seems to potentially contact *CDC6*.

**Table 3.** Additional ALS saDMP gene annotations identified by chromatin loop analysis.

| CpG | Chromosome | Start Position | End Position | Strand | Anchor One | Anchor Two | Effect |
| --- | --- | --- | --- | --- | --- | --- | --- |
| cg12393318 | chr5 | 87975817 | 150829561 | * | *LINC00461* | *SLC36A1* | Female biased |
| cg23094674 | chr1 | 204329128 | 2131439 | * | *LINC00628* | *FAAP20* | Female biased |
| cg01681881 | chr11 | 57397473 | 102160731 | * | NA | NA | Female biased |
| cg15763020 | chr2 | 38967142 | 11840126 | * | *SRSF7* | *LPIN1* | Female biased |
| cg00480331 | chr3 | 177067788 | 187457788 | * | NA | *BCL6* | Female biased |
| cg03053374 | chr17 | 40552018 | 38466252 | * | *CAVIN1* | *CDC6* | Female biased |
| cg04183381 | chr22-21 | 30775989 | 15782321 | * | *RNF215* | NA | Male biased |
| cg03746976 | chr16 | 58033904 | 67583903 | * | *USB1* | *RIPOR1* | Male biased |
| cg09409484 | chr1 | 45765672 | 54875673 | * | *LINC01144* | *SSBP3* | Male biased |
| cg05335422 | chr12 | 51733784 | 65003780 | * | *CELA1* | NA | Male biased |
| cg21585707 | chr7 | 25129619 | 5539631 | * | NA | *FBXL18* | Male biased |
| cg21585707 | chr7 | 101921951 | 5539631 | * | *CUX1* | *FBXL18* | Male biased |
| cg11741201 | chr11 | 64077472 | 35641548 | * | *TRMT112* | *FJX1* | Male biased |
| cg20813374 | chr6 | 44207737 | 35657777 | * | *HSP90AB1* | *FKBP5* | Male biased |
| cg08234504 | chr5 | 67755827 | 139019585 | * | NA | NA | Male biased |

### Enrichment of Transcription Factor Motifs at ALS saDMPs

Recognizing the critical role of enhancers in transcriptional regulation, we investigated whether differential methylation at these sites could alter transcription factorbinding dynamics. To this end, we performed a transcription factor motif enrichment analysis at the ALS saDMPs. This analysis aimed to identify specific transcription factor motifs that were overrepresented, thereby suggesting potential TFs that might interact with these enhancers. Through this analysis, we identified two enriched transcription factor motifs in the ALS saDMPs. The ESR1 motif, encoding the estrogen receptor alpha, was significantly enriched at ALS saDMPs. ESR1 is a critical regulator of estrogen-responsive genes and has been implicated in various cellular processes, including neuronal growth, differentiation, and survival. The enrichment of ESR1 motifs suggests that estrogen signaling pathways may be dysregulated in ALS, potentially contributing to disease progression. Additionally, the REST (RE1-Silencing Transcription factor) motif was found to be enriched at these ALS saDMPs. REST is a well-known regulator of a genetic network involved in neurodegeneration. It acts as a transcriptional repressor, silencing neuronal genes in non-neuronal tissues and modulating neurogenesis.

### Survival Outcomes Linked to Methylation at Two ALS saDMPs

To investigate the relationship between DNA methylation at specific CpG sites and overall survival in ALS, we conducted a Cox proportional hazards analysis. We focused on the ALS saDMPs that were identified as significant in our meta-analysis. We used false discovery rate (FDR)-adjusted p-values in the Cox proportional hazards model to account for multiple testing across the 226 CpG sites, while adjusting for age, age of onset, C9 status, age at diagnosis, sex, and cell type composition. Our analysis revealed that two ALS saDMPs (cg14380013 and cg06729676) showed a significant association with survival, indicating that not only do these sites show a sex difference but also are associated with survival in ALS (Figure 3). The results of these survival analyses have been preserved in a series of Cox proportional hazards models, which were filtered to retain only those with statistically significant associations. Specifically, cg14380013 is located at the *UNC119B* gene, a transport factor which plays a role in the pathogenesis of ALS, and cg06729676 is located at the NF1 gene.

**Figure 3.**
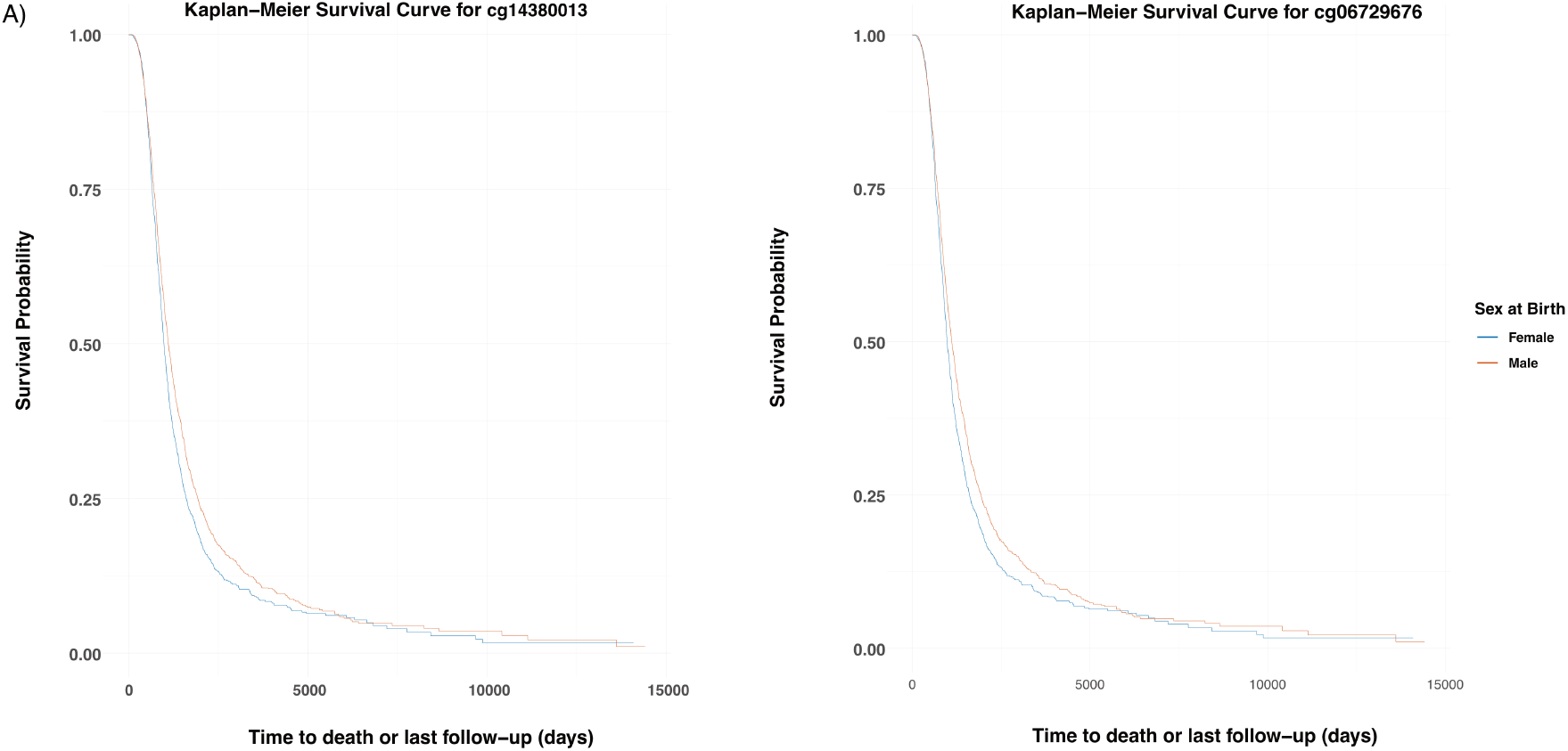
Kaplan-Meier survival curves for two CpG sites stratified by sex.(A) Kaplan-Meier survival curve for cg14380013, and (B) Kaplan-Meier survival curve for cg06729676. Survival probability is plotted on the y-axis, while time to death or last follow-up (days) is on the x-axis. The curves are stratified by sex at birth, with females shown in blue and males shown in orange. Survival was measured based on time to death and survival status.

In addition to the Cox regression analysis, we further investigated potential differences in survival between sexes by plotting Kaplan-Meier survival curves stratified by sex at birth. This allowed us to visually assess whether there are observable differences in survival probabilities between males and females for the two CpG sites identified.

As shown in Figure 3, the Kaplan-Meier curves for both cg14380013 (A) and cg06729676 (B) exhibit very similar patterns between males and females. The survival curves for males (orange) and females (blue) are nearly overlapping, indicating minimal to no observable differences in survival between the sexes for these CpG sites. Both curves follow a typical survival decline, suggesting that while the methylation levels at these CpG sites are associated with overall survival, sex does not appear to substantially influence the survival probability in these cases.

## Methods

### Datasets

This study aimed to identify sex-biased differential DNA methylation in individuals diagnosed with definite, probable, and probable laboratory-supported ALS according to the revised El Escorial Criteria^26^. Whole blood DNA methylation data were collated from individuals involved in the Project MinE sequencing consortium across 14 countries, employing a 2:1 case/control ratio. Population-based controls were matched for age, sex, and geographical region in a 1:2 ratio, without screening for (subclinical) signs of ALS. Experimental batches were processed uniformly in the same laboratory, resulting in 44 independent batches after quality control.

We applied extensive quality control measures, leading to the exclusion of 756 (7.2%) individuals (based on several technical metrics, relatedness, genotype concordance, and sex concordance) and 175,134 (24%) probes (based on technical metrics, cross-reactivity, and overlap with common single-nucleotide polymorphisms). The investigators were not blinded to the experimental conditions during experiments and analyses. Following these quality checks, our final dataset consisted of 9274 participants (5529 males and 3975 females)

### DNA Methylation Data

Sample collection has been previously described in^12^. Analyses were stratified by array technology (MinE 450k and MinE EPIC) and external Australian data were split into two strata based on signal intensities (AUS1 and AUS2) as previously reported^12^.

To account for the heterogeneous nature of whole blood, where varying cell type proportions can confound analyses, we employed the Houseman method^27^ to estimate cell type composition using DNA methylation signatures. The estimateCellCounts function within the bigmelon package^28^ was used to estimate the relative proportions of several key cell types, including Granulocytes, Mononuclear cells, Natural Killer cells, CD4+ T cells, CD8+ T cells, and B cells for all individuals.

Furthermore, to determine whether observed sex differences were independent of age, we performed a Mann–Whitney U test to compare age distributions between male and female individuals. The test results indicated no statistically significant difference in age distribution (p value 0.07; median ages of 62 and 63 for males and females, respectively), supporting the notion that our observed sex differences are not influenced by age disparities.

### Identifying sex-associated autosomal differential methylation

Sex-associated autosomal differentially methylated positions (saDMPs) were identified by performing linear modelling using the limma package in R [92] using ALS status, sex annotation and Beta values while adjusting for age, cell type proportions, principal components and batch effects. Correction for multiple testing was performed with the Benjamini–Hochberg false discovery rate method (FDR values). We further used the Bayesian method for controlling p value inflation using the R package bacon for each of our data sets^29^. We performed individual epigenome wide association studies in 4 cohorts. (I) Project MiNE samples based on the 450k array (II) Project MiNE samples based on the EPIC array (III) Australian strata 1 and (III) Australian strata 2. These were then meta analysed, which is described below. To ensure the validity of genome-wide discoveries, controlling the false positive rate is crucial. Traditionally, genomic inflation factor (lambda) has been employed to quantify inflation in genome-wide association studies (GWAS) of genetic variants. However, in epigenome-wide association studies (EWAS), where small effects from numerous CpGs may be associated with the phenotype, lambda can overestimate test-statistic inflation. To address this, Iterson et al. developed a Bayesian method implemented in the Bioconductor package bacon, which estimates inflation in EWAS based on empirical null distributions^29^. In this study, we utilized quantile–quantile (QQ) plots and estimated genomic inflation factors using both conventional methods and the bacon approach. The bacon method was further employed to derive inflation-corrected effect sizes, standard errors, and P-values for each cohort, facilitating meta-analysis implemented in the R package metafor^30^.

### Meta analysis

A meta-analysis was undertaken using the ‘metafor’ package in R to combine data from four separate cohorts: 450k, Epic Array, Aus Strata 1, and Aus Strata 2. Meta-analysis is a tool that aggregates findings from multiple studies to yield more robust conclusions than individual studies alone. The ‘metafor’ package allows for the incorporation of various study characteristics, such as sample size and effect size estimates, while addressing between-study differences. In this analysis, fixed effects models were employed to accommodate potential variability across the cohorts. These models assume that the true effect size remains constant across studies, while still capturing within-study variability. The determination of the overall effect size and its confidence interval relied on utilizing the standard errors of the effect size estimates from each cohort. These standard errors quantify the uncertainty associated with the effect size estimates and are pivotal for appropriately weighting each study in the meta-analysis. By incorporating standard errors, the meta-analysis effectively considers both the precision of the effect size estimates and the sample sizes of the cohorts. In essence, the meta-analysis facilitated by the ‘metafor’ package facilitated the integration of data from multiple cohorts, addressing between-study variability and providing a comprehensive understanding of the collective evidence across the included studies.

### Genomic annotation of CpG sites

The method used for annotating CpG sites on the 450k and EPIC array has previously been described by^31^. However, briefly we annotated the ALS saDMPs to CpG islands, CpG shores, CpG shelves and open sea regions using the manufacturer supplied annotation data (specifically the MethylationEPICv 10B2 manifest]file). Furter, we annotated them to additional regions in the genome such as enhancers, promoters and more using data from UCSC genome browser.

### Gene ontology analyses

To explore the functional implications of significant CpGs, we conducted Gene Ontology (GO) and Kyoto Encyclopedia of Genes and Genomes (KEGG) analyses. These analyses were performed using the gometh function within the missMethyl package^32^.The gometh function is specifically designed to assess gene ontology enrichment while considering the variable number of probes per gene present on the EPIC array. This is crucial as the EPIC array may contain multiple probes targeting the same gene, and accounting for this variability ensures accurate interpretation of enrichment results.

GO analysis provides insights into the biological processes, molecular functions, and cellular components associated with the identified CpGs. On the other hand, KEGG analysis offers information on the pathways in which these CpGs are involved, shedding light on potential molecular mechanisms underlying observed associations. By incorporating both GO and KEGG analyses, we aim to comprehensively elucidate the functional relevance of the significant CpGs identified in our study, providing valuable insights into their potential roles in biological processes and pathways.

### Enrichment of saDMPs in transcription factor binding motifs

To elucidate potential regulatory mechanisms underlying sex-associated differentially methylated positions (saDMPs), we conducted an enrichment analysis of known motifs using the R package PWMEnrich^33^. This analysis leveraged the MotifDb collection of transcription factor (TF) motifs [99]. Specifically, we utilized the getSeq function and constructed ‘dnastringsets’ to extract DNA sequences within a 50 bp range surrounding the saDMPs identified in amyotrophic lateral sclerosis (ALS) samples. These sequences were then compared against a human background to unveil enriched motifs. A motif was considered enriched if its adjusted p-value was smaller than 0.05, indicating statistical significance. By identifying enriched transcription factor binding motifs associated with saDMPs, we aimed to gain insights into the potential regulatory roles of these CpG sites in gene expression regulation. Integration of these findings with gene expression data provides a comprehensive understanding of how DNA methylation alterations at specific loci may influence transcription factor binding and subsequent gene expression changes in the context of ALS and sex differences.

### Overlap of saDMP’s with chromatin loops

We investigated the potential interactions between ALS saDMPs and distal genes in three-dimensional space using Hi-C data available from the Gene Expression Omnibus (GEO) under accession number GSE124974 for white blood cells and neutrophils. The Hi-C library preparation was conducted utilizing the Arima-HiC kit, and preprocessing of the data was executed employing Juicer command line tools^34^. Initially, reads were aligned to the human genome (hg38) using BWA-mem. Subsequently, the Juicer pre-processing pipeline was employed for further data processing. To identify chromatin loops, we employed the HICCUPS tool from Juicer, utilizing a resolution of 10 kilobases (Kb). Following loop calling, we constructed GenomicInteractions objects to annotate sex-associated DMPs (saDMPs) to loop anchors. This annotation was performed using the findOverlaps function from the GenomicRanges package. Subsequently, we annotated the corresponding loop anchor to the relevant gene identifiers (IDs). This comprehensive analysis allowed us to explore potential long-range interactions between DMPs and distal genes, providing insights into the regulatory landscape of these genomic regions in the context of white blood cells and neutrophils.

### Survival analysis and Kaplan-Meier survival curves

To investigate the association between DNA methylation and overall survival in ALS, we first identified ALS saDMPs associated with survival using Cox proportional hazards models. The Cox models were adjusted for potential confounders including age, sex, age at diagnosis, age at onset, C9orf72 repeat expansion status, and cell type proportions. Specifically, we focused on the 226 CpG sites that were significant in our meta-analysis. We used false discovery rate (FDR)-adjusted p-values to account for multiple testing across these CpG sites, ensuring robust significance thresholds. We employed the coxph function from the survival package in R for this analysis. To further explore potential sex differences in the association between methylation and survival, Kaplan-Meier survival curves were plotted. We stratified these survival curves by sex at birth. The survival curves were generated using the survfit function in the survival package, and plotted using ggsurvplot from the survminer package.

The Kaplan-Meier plots allowed us to visually assess and compare survival probabilities over time between males and females for the identified CpG sites. The log-rank test was used to statistically evaluate differences between the survival curves of males and females. This approach enabled us to determine whether the survival trajectories diverged significantly based on sex. All analyses were conducted in R, and results were considered statistically significant at a threshold of FDR-adjusted p < 0.05.

## Discussion

Here, we conducted a comprehensive meta-analysis to investigate sex-specific differences in DNA methylation among individuals with amyotrophic lateral sclerosis (ALS). We identified 226 ALS-Sex associated DMPs (ALS-saDMPs). Although the differences in methylation levels between males and females were relatively small as indicated by the small effect sizes, they were statistically significant. Notably, 54% of these CpGs were more highly methylated in females, while 46% were more methylated in males. This subtle yet significant difference suggests that sex-specific methylation patterns exist, and while their overall impact on gene expression and disease mechanisms may be limited, these changes could still be biologically relevant, particularly if they occur in regulatory regions controlling the expression of genes crucial for neuronal health.

However, there may be a few other reasons for the small effect sizes, for example studying DNA methylation changes in ALS using whole blood may provide limited insights compared to more relevant tissues, such as brain tissue, where the disease predominantly manifests. However, the collection of brain tissue for such studies presents significant logistical and ethical challenges and so performing studies of this nature using more accessible tissues like whole blood can still be insightful. Additionally, the coverage of CpG sites on the 450k and EPIC arrays may not be sufficient to fully address our research hypothesis. For instance, array-based methods like the Illumina 450k array focus on a selected set of CpG sites deemed functionally important, which may not encompass all relevant regions of the genome. This could limit the ability to detect significant associations or fully understand the complex relationship between DNA methylation and ALS. Furthermore, even when statistically significant, the effect sizes of DNA methylation changes are often small, complicating the interpretation of their biological relevance. Thus, while DNA methylation offers valuable insights, it may not always capture the complete picture of disease mechanisms or predict phenotypic outcomes effectively.

Nevertheless, despite the small effect sizes observed, the fact that many differentially methylated positions (DMPs) overlap with known ALS-associated genes or have been previously implicated in similar studies lends credibility to our findings. For instance, HLA-F emerged as a top hit, containing three ALS-associated DMPs in total, with 2 of these being among the top 10 most significant and they were all male biased. HLA-F is known for its protective role in ALS, as evidenced by its reduced expression in motor neurons in ALS cases and its ability to extend survival in animal models through increased expression^35^. Furthermore, the HLA-F loci was also identified as a male-specific risk loci in a recent sex specific GWAS of ALS^36^ which may point towards a possible interplay between genetic and epigenetic factors in the context of sex differences of ALS.

Furthermore, the top hit was at cg16904330 located in the *ADPRH* and was female biased, meaning it was more highly methylated in females. *ADPRH* has previously been linked specifically to genetic risk of ALS^37,38^. Interestingly, *KCNQ1* contained two ALS saDMPs, one of which was male-biased and the other female-biased. *KCNQ1* has recently been reported as a potentially significant target for managing hyperexcitability in motor neurons, a pathological feature that can exacerbate neurodegeneration. Notably, KCNQ1 is an imprinted gene, meaning that it is expressed in a parent-of-origin-specific manner^39^, which adds an additional layer of complexity to understanding its role in ALS. The sex-biased methylation patterns observed in KCNQ1 may interact with imprinting mechanisms, possibly influencing allele-specific expression differently in males and females. This could have important implications for how hyperexcitability and neurodegeneration are modulated in a sex-specific manner in ALS.

Additionally, other top hits were also found in genes that have previously been linked to ALS such as *SERPINB6, MIR130A, TRAF31P2 and UBASH3B*^25,40,41^. Further, the analysis of the genomic locations of these ALS-associated sex-differentially methylated positions (saDMPs) revealed an enrichment in CpG shores, with a slight enrichment at CpG islands, and a depletion in open sea regions, this is consistent with previous research looking at sex differences of DNA methylation in whole blood^31^. Methylation in CpG islands typically functions to serve long term silencing of genes and CpG islands are often unmethylated, especially when located at transcription start sites. CpG shores are adjacent to CpG islands and methylation in these regions is also highly correlated with gene silencing^42^. This distribution suggests a possible functional relevance, as CpG shore and island methylation is often involved in gene regulation. Further annotation of these saDMPs to genomic elements, such as enhancers and 5’UTRs, indicated a significant enrichment in enhancers—regions that are critical for gene expression, often through long-range chromatin interactions. This enrichment highlights the potential for these methylation changes to exert influence over genes located far from the DMPs themselves, adding another layer of complexity to the gene regulation mechanisms at play in ALS.

Adding to this idea, it’s important to note that gene expression is also influenced by chromatin organisation and 3D interactions, mentioned above. This suggests that the interplay between epigenetic changes and 3D genome organisation may work together to explain some of the sex biases seen in diseases like ALS. In line with this, previous research has highlighted that sex specific chromatin accessibility is associated with sex specific gene expression^43^. Further, it has also been shown that 3D genome architecture can affect sex-biased gene expression through both direct and indirect effects of cohesin and CTCF looping, which influence enhancer interactions with sex-biased genes^44^. Therefore, given that it is well established that DNA methylation can impact genome organisation^45–48^ it is possible that sex-specific methylation patterns may interact with chromatin remodeling processes, influencing 3D genome architecture. This interaction could drive differences in gene expression between sexes, potentially contributing to the sex biases observed in diseases like ALS. Our exploration of this idea of the presence of potential chromatin loops involving ALS saDMPs provided additional insights into their functional implications. We identified saDMPs that potentially contact promoter regions of known ALS genes through chromatin loops, suggesting that these methylation changes could impact gene expression by altering the three-dimensional structure of the genome. For instance, a female-biased saDMP on chromosome 5 was found to potentially contact the promoter region of the *SLC36A1* gene, which has been implicated in ALS and Alzheimer’s disease^49^. Further, a male biased saDMPs on chromosome 6 (cg20813374) located at gene *FKBP5* was found to contact the gene *HSP90AB1*. Interestingly, *HSP90AB1* encodes for a heat shock protein involved in protein folding and stability, and has been identified as a substrate of Cyclin-F, suggesting that its dysregulation through ubiquitination may contribute to ALS pathogenesis by impairing proteostasis^50^.

To elucidate the potential regulatory mechanisms underlying the identified saDMPs, we performed a transcription factor motif enrichment analysis. Two enriched transcription factormotifs were identified: ESR1, encoding the estrogen receptor alpha, and REST, a well-known regulator of neurodegeneration-related genes. The enrichment of ESR1 motifs suggests that estrogen signaling pathways may be dysregulated in ALS, potentially contributing to disease progression. This finding is supported by previous studies showing the neuroprotective effects of estrogen in models of neurodegeneration. The enrichment of REST motifs indicates that methylation changes could influence REST binding, thereby affecting the expression of genes involved in ALS.

Despite these significant findings, our study has some limitations. The relatively small differences in methylation levels between sexes, although statistically significant, may suggest that other genetic and environmental factors also play crucial roles in ALS pathogenesis. Furthermore, the lack of significant enriched terms in GO and KEGG pathway analyses indicates that the identified saDMPs do not prominently cluster within particular biological pathways or processes. This diffuse pattern of methylation changes suggests a more complex interplay of genetic and epigenetic factors in ALS, where sex differences in methylation are part of a larger, multifactorial context influencing disease development and progression.

Understanding sex-specific DNA methylation changes in ALS is crucial for several reasons. First, it could elucidate the mechanisms underlying the observed differences in disease manifestation and progression between men and women. Second, it might reveal novel biomarkers for ALS that are specific to each sex, thereby improving diagnostic accuracy and prognostic predictions. Lastly, this knowledge could inform the development of sex-specific therapeutic strategies, potentially enhancing the effectiveness of treatments.

In conclusion, our study provides valuable insights into the sex-specific epigenetic landscape of ALS. The identification of sex-biased differential DNA methylation patterns and their potential functional implications highlights the importance of considering sex as a biological variable in ALS research. Future studies should further explore the interplay between genetic, epigenetic, and environmental factors in ALS to develop more personalized and effective interventions for this debilitating disease. Additionally, future work should prioritize the investigation of more relevant tissues, such as the brain or central nervous system, where ALS pathology originates. Examining these tissues may provide deeper insights into the molecular mechanisms driving ALS and improve our understanding of sex-specific disease progression and therapeutic targets.

## Acknowledgements

Samples used in this research were in part obtained from the UK National DNA Bank for MND Research, funded by the MND Association and the Wellcome Trust. We thank people with MND and their families for their participation in this project. We acknowledge sample management undertaken by Biobanking Solutions funded by the Medical Research Council at the Centre for Integrated Genomic Medical Research, University of Manchester.

The authors acknowledge use of the research computing facility at King’s College London, Rosalind (https://rosalind.kcl.ac.uk), which is delivered in partnership with the National Institute for Health Research (NIHR) Biomedical Research Centres at South London & Maudsley and Guy’s & St Thomas’ NHS Foundation Trusts, and part-funded by capital equipment grants from the Maudsley Charity (award 980) and Guy’s & St Thomas’ Charity (TR130505). The views expressed are those of the author(s) and not necessarily those of the NHS, the NIHR, King’s College London, or the Department of Health and Social Care.

The authors acknowledge the National Institute for Health Research (NIHR) Biomedical Research Centre at South London & Maudsley NHS Foundation Trust and King’s College London. The authors also acknowledge Health Data Research UK, which is funded by the UK Medical Research Council, Engineering and Physical Sciences Research Council, Economic and Social Research Council, Department of Health and Social Care (England), Chief Scientist Office of the Scottish Government Health and Social Care Directorates, Health and Social Care Research and Development Division (Welsh Government), Public Health Agency (Northern Ireland), British Heart Foundation, and Wellcome Trust.

## Supplementary Information

### Full List of ALS Sex-Associated Differentially Methylated Positions (saDMPs)

The complete list of ALS-associated sex-differentially methylated positions (ALS saDMPs) identified in this study is provided in the supplementary files. These include all 226 CpG sites that passed significance thresholds in our meta-analysis, annotated to 159 unique genes. For each saDMP, detailed information is provided on its genomic location, direction of bias, annotated gene, and significance levels (FDR and Bonferroni).

This supplementary information serves as an extended resource, allowing interested readers to access the full dataset underpinning the findings discussed in the main text.

## Supplementary Figures and Tables

**Supplementary Figure 1.**
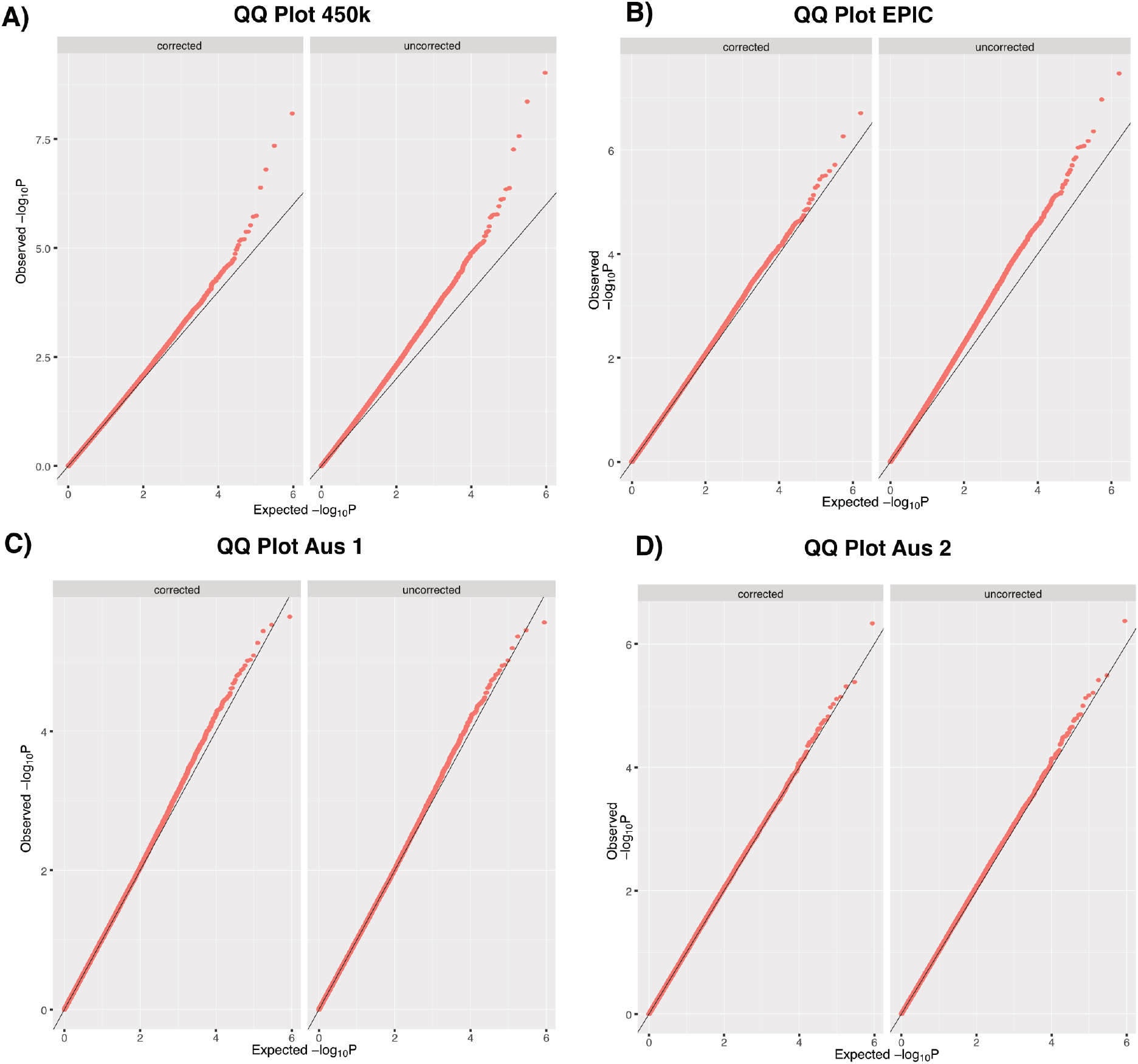
Quantile-Quantile (QQ) plots and lambda values showing the distribution of adjusted p-values against the null distribution for EWAS of sex differences in DNA methylation associated with ALS. The genomic inflation lambda score is indicated in each QQ plot to show statistical inflation of p-values. Each panel includes both corrected (left) and uncorrected (right) QQ plots to compare the effects of BACON correction on the distribution of p-values, with the black diagonal line representing the expected line under the null hypothesis. (A) 450k data, (B) EPIC data, (C) Aus 1 data, (D) Aus 2 data.

**Supplementary Figure 2.**
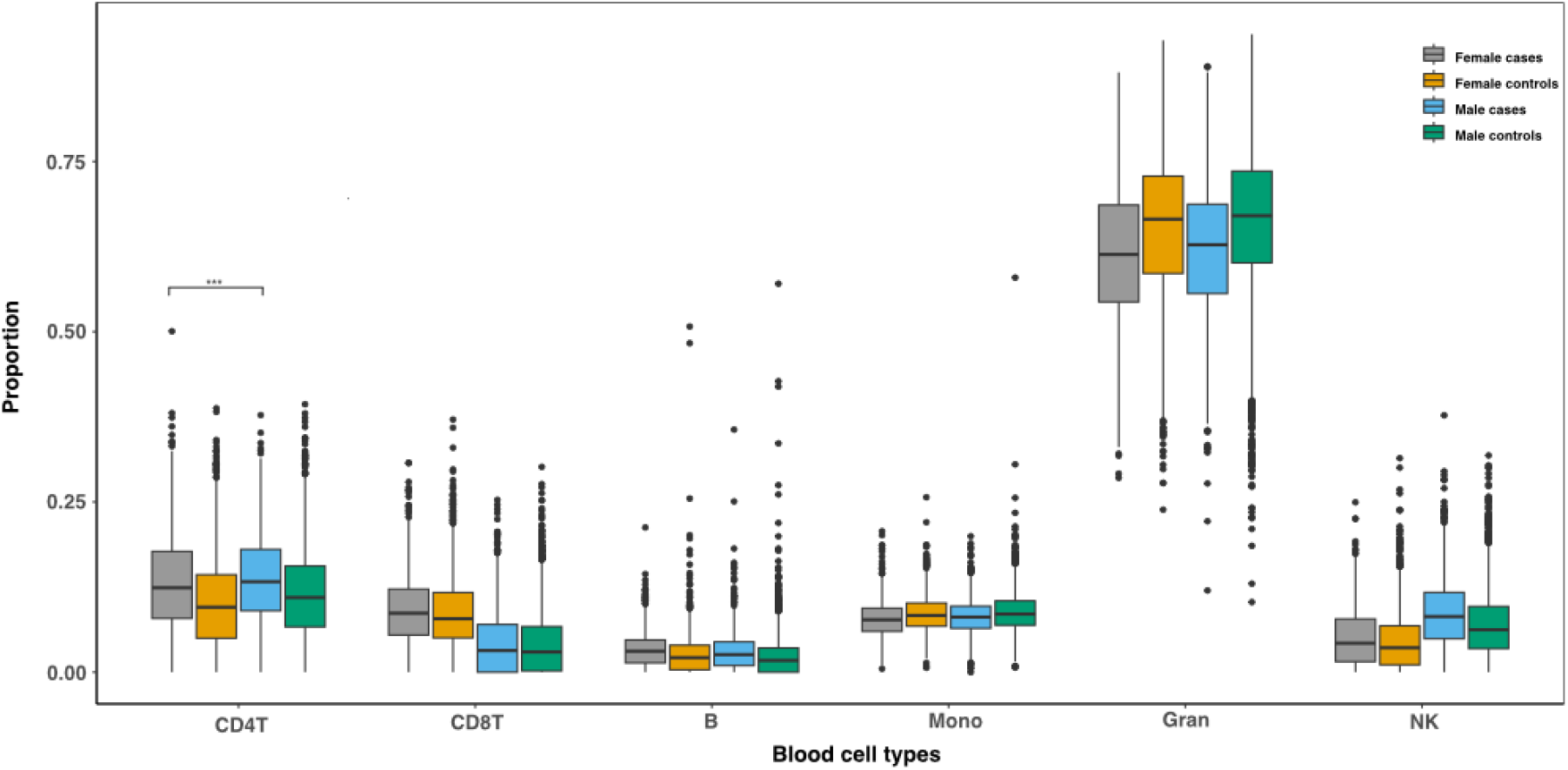
Distribution of blood cell type proportions between males and females in cases and controls. The box plots show the proportions of various blood cell types (CD4T, CD8T, B, Mono, Gran, NK) across four groups: female cases (orange), female controls (gray), male cases (blue), and male controls (green). Each box plot displays the median (central line in the box), the interquartile range (IQR; bounds of the box), and outliers (individual points outside the whiskers). The whiskers extend to the furthest data point within 1.5 times the IQR from the box. Notably, there is a significant difference in the proportions of CD4T cells between female cases and female controls, as indicated by asterisks (^***^ p < 0.001).

## References

1. Van Es, M. A. et al. Amyotrophic lateral sclerosis. The Lancet 390, 2084–2098 (2017).

2. Siddique, T. & Ajroud-Driss, S. Familial amyotrophic lateral sclerosis, a historical perspective. Acta Myol. 30, 117 (2011).

3. Chen, S., Sayana, P., Zhang, X. & Le, W. Genetics of amyotrophic lateral sclerosis: an update. Mol. neurodegeneration 8, 1–15 (2013).

4. Wang, H., Guan, L. & Deng, M. Recent progress of the genetics of amyotrophic lateral sclerosis and challenges of gene therapy. Front. Neurosci. 17, 1170996 (2023).

5. Kamel, F., Umbach, D. M., Munsat, T. L., Shefner, J. M. & Sandler, D. P. Association of cigarette smoking with amyotrophic lateral sclerosis. Neuroepidemiology 18, 194–202 (1999).

6. Menounos, S., Hansbro, P. M., Diwan, A. D. & Das, A. Pathophysiological correlation between cigarette smoking and amyotrophic lateral sclerosis. NeuroSci 2, 120–134 (2021).

7. Fang, F. et al. Workplace exposures and the risk of amyotrophic lateral sclerosis. Environ. Heal. Perspectives 117, 1387–1392 (2009).

8. Andrew, A. S. et al. Environmental and occupational exposures and amyotrophic lateral sclerosis in new england. Neurodegener. Dis. 17, 110–116 (2017).

9. Wu, F. et al. Exposure to ambient air toxicants and the risk of amyotrophic lateral sclerosis (als): A matched case control study. Environ. Res. 242, 117719 (2024).

10. Gibney, E. & Nolan, C. Epigenetics and gene expression. Heredity 105, 4–13 (2010).

11. Singal, R. & Ginder, G. D. Dna methylation. Blood, The J. Am. Soc. Hematol. 93, 4059–4070 (1999).

12. Hop, P. J. et al. Genome-wide study of dna methylation shows alterations in metabolic, inflammatory, and cholesterol pathways in als. Sci. translational medicine 14, eabj0264 (2022).

13. Zhang, M. et al. Combined epigenetic/genetic study identified an als age of onset modifier. Acta Neuropathol. Commun. 9, 75 (2021).

14. Bennett, S. A., Tanaz, R., Cobos, S. N. & Torrente, M. P. Epigenetics in amyotrophic lateral sclerosis: a role for histone post-translational modifications in neurodegenerative disease. Transl. research 204, 19–30 (2019).

15. Young, P. E., Kum Jew, S., Buckland, M. E., Pamphlett, R. & Suter, C. M. Epigenetic differences between monozygotic twins discordant for amyotrophic lateral sclerosis (als) provide clues to disease pathogenesis. PLoS One 12, e0182638 (2017).

16. Cai, Z., Jia, X., Liu, M., Yang, X. & Cui, L. Epigenome-wide dna methylation study of whole blood in patients with sporadic amyotrophic lateral sclerosis. Chin. Med. J. 135, 1466–1473 (2022).

17. Coppedè, F. et al. Increase in dna methylation in patients with amyotrophic lateral sclerosis carriers of not fully penetrant sod1 mutations. Amyotroph. Lateral Scler. Frontotemporal Degener. 19, 93–101 (2018).

18. Yazar, V. et al. Dna methylation analysis in monozygotic twins discordant for als in blood cells. Epigenetics Insights 16, 25168657231172159 (2023).

19. Wijesekera, L. C. & Nigel Leigh, P. Amyotrophic lateral sclerosis. Orphanet journal rare diseases 4, 1–22 (2009).

20. Manjaly, Z. R. et al. The sex ratio in amyotrophic lateral sclerosis: A population based study. Amyotroph. Lateral Scler. 11, 439–442 (2010).

21. Trojsi, F., D’Alvano, G., Bonavita, S. & Tedeschi, G. Genetics and sex in the pathogenesis of amyotrophic lateral sclerosis (als): is there a link? Int. journal molecular sciences 21, 3647 (2020).

22. Moore, L. D., Le, T. & Fan, G. Dna methylation and its basic function. Neuropsychopharmacology 38, 23–38 (2013).

23. Huang, X. et al. Human amyotrophic lateral sclerosis excitability phenotype screen: Target discovery and validation. Cell reports 35 (2021).

24. Dulberger, C. L. et al. Human leukocyte antigen f presents peptides and regulates immunity through interactions with nk cell receptors. Immunity 46, 1018–1029 (2017).

25. Liguori, M. et al. Dysregulation of micrornas and target genes networks in peripheral blood of patients with sporadic amyotrophic lateral sclerosis. Front. molecular neuroscience 11, 288 (2018).

26. Brooks, B. R., Miller, R. G., Swash, M. & Munsat, T. L. El escorial revisited: revised criteria for the diagnosis of amyotrophic lateral sclerosis. Amyotroph. lateral sclerosis other motor neuron disorders 1, 293–299 (2000).

27. Houseman, E. A. et al. Dna methylation arrays as surrogate measures of cell mixture distribution. BMC bioinformatics 13, 1–16 (2012).

28. Gorrie-Stone, T. J. et al. Bigmelon: tools for analysing large dna methylation datasets. Bioinformatics 35, 981–986 (2019).

29. van Iterson, M., van Zwet, E. W., Consortium, B. & Heijmans, B. T. Controlling bias and inflation in epigenome-and transcriptome-wide association studies using the empirical null distribution. Genome biology 18, 1–13 (2017).

30. Viechtbauer, W. Conducting meta-analyses in R with the metafor package. J. Stat. Softw. 36, 1–48, DOI: 10.18637/jss.v036.i03 (2010).

31. Grant, O. A., Wang, Y., Kumari, M., Zabet, N. R. & Schalkwyk, L. Characterising sex differences of autosomal dna methylation in whole blood using the illumina epic array. Clin. epigenetics 14, 62 (2022).

32. Phipson, B. & Maksimovic, J. missmethyl: Analysing illumina humanmethylation450 beadchip data.. Stojnić, R. Overview of the pwmenrich package. (2019).

33. Durand, N. C. et al. Juicer provides a one-click system for analyzing loop-resolution hi-c experiments. Cell systems 3, 95–98 (2016).

34. Song, S. et al. Major histocompatibility complex class i molecules protect motor neurons from astrocyte-induced toxicity in amyotrophic lateral sclerosis. Nat. medicine 22, 397–403 (2016).

35. Byrne, R. P. et al. Sex-specific risk loci and modified mef2c expression in als. medRxiv 2024–05 (2024).

36. Szymanski, J. Genetic analysis of familial alzheimer’s disease, primary lateral sclerosis and paroxysmal kinesigenic dyskinesia: a tool to uncover common mechanistic points. (2021).

37. Annan, L. V. N. D. The pathophysiology of Spinal Bulbar Muscular Atrophy: a longitudinal analysis of muscle and spinal cord. Ph.D. thesis, UCL (University College London) (2017).

38. Lewis, A. et al. Epigenetic dynamics of the kcnq1 imprinted domain in the early embryo. (2006).

39. Noori, A., Mezlini, A. M., Hyman, B. T., Serrano-Pozo, A. & Das, S. Systematic review and meta-analysis of human transcriptomics reveals neuroinflammation, deficient energy metabolism, and proteostasis failure across neurodegeneration. Neurobiol. disease 149, 105225 (2021).

40. Mamoor, S. Differential expression of traf3ip2 in amyotrophic lateral sclerosis., DOI: 10.31219/osf.io/an4hj (2022).

41. Irizarry, R. A. et al. The human colon cancer methylome shows similar hypo-and hypermethylation at conserved tissue-specific cpg island shores. Nat. genetics 41, 178–186 (2009).

42. Kukurba, K. R. et al. Impact of the x chromosome and sex on regulatory variation. Genome research 26, 768–777 (2016).

43. Werner, R. J. et al. Sex chromosomes drive gene expression and regulatory dimorphisms in mouse embryonic stem cells. Biol. sex differences 8, 1–18 (2017).

44. Monteagudo-Sánchez, A., Noordermeer, D. & Greenberg, M. V. The impact of dna methylation on ctcf-mediated 3d genome organization. Nat. Struct. & Mol. Biol. 31, 404–412 (2024).

45. Roseman, S. A. et al. Dna methylation insulates genic regions from ctcf loops near nuclear speckles. bioRxiv 2023–07 (2023).

46. Damaschke, N. A. et al. Ctcf loss mediates unique dna hypermethylation landscapes in human cancers. Clin. Epigenetics 12, 1–13 (2020).

47. Soochit, W. et al. Ctcf chromatin residence time controls three-dimensional genome organization, gene expression and dna methylation in pluripotent cells. Nat. cell biology 23, 881–893 (2021).

48. Wainberg, M., Andrews, S. J. & Tripathy, S. J. Shared genetic risk loci between alzheimer’s disease and related dementias, parkinson’s disease, and amyotrophic lateral sclerosis. Alzheimer’s Res. & Ther. 15, 113 (2023).

49. Siebert, A., Gattringer, V., Weishaupt, J. H. & Behrends, C. Als-linked loss of cyclin-f function affects hsp90. Life Sci. Alliance 5 (2022).

